# Slow-wave brain connectivity predicts executive functioning and group belonging in socially vulnerable individuals

**DOI:** 10.1101/2023.07.19.549808

**Authors:** Renzo C. Lanfranco, Fabienne dos Santos Sousa, Pierre Musa Wessel, Álvaro Rivera-Rei, Tristán A. Bekinschtein, Boris Lucero, Andrés Canales-Johnson, David Huepe

## Abstract

Important efforts have been made to describe the neural and cognitive features of healthy and clinical populations. However, the neural and cognitive features of socially vulnerable individuals remain largely unexplored, despite their proneness to developing neurocognitive disorders. Socially vulnerable individuals can be characterised as socially deprived, having a low socioeconomic status, suffering from chronic social stress, and exhibiting poor social adaptation. While it is known that such individuals are likely to perform worse than their peers on executive function tasks, studies on healthy but socially vulnerable groups are lacking. In the current study, we explore whether neural power and connectivity signatures can characterise executive function performance in healthy but socially vulnerable individuals, shedding light on the impairing effects that chronic stress and social disadvantages have on cognition. We measured resting-state electroencephalography and executive functioning in 38 socially vulnerable participants and 38 matched control participants. Our findings indicate that while neural power was uninformative, lower delta and theta phase synchrony are associated with worse executive function performance in all participants, whereas delta phase synchrony is higher in the socially vulnerable group compared to the control group. Finally, we found that delta phase synchrony and years of schooling are the best predictors for belonging to the socially vulnerable group. Overall, these findings suggest that exposure to chronic stress due to socioeconomic factors and a lack of education are associated with changes in slow-wave neural connectivity and executive functioning.

## 1. INTRODUCTION

Social and economic factors can play a crucial role in people’s mental well-being by easing or hampering their access to education, healthcare, social security, and work opportunities. Socioeconomically vulnerable individuals typically struggle at getting access to such resources, which may lead to chronic stress, oftentimes precluding them from a healthy cognitive development (Cermakova et al., 2018; Migeot et al., 2022). Indeed, they frequently experience a higher rate of domestic problems and live in areas with higher crime and drug abuse rates, and scarce recreational spaces (De Nadai et al., 2020; Engelberg et al., 2016; Giles-Corti & Donovan, 2002). Many studies have thoroughly characterised cognitive functioning in groups that present risk factors for developing psychiatric disorders (i.e., clinical populations), finding a generalisable decrease in their executive function performance (Romer & Pizzagalli, 2021; Testa & Pantelis, 2009). However, studies characterising cognitive functioning in healthy but socially vulnerable groups are still lacking, even though such groups are particularly at risk of chronic stress and mental illness (Baum et al., 1999).

Executive function encompasses a set of cognitive skills crucial for the control of behaviour, including focusing attention, planning, remembering relevant recent information, thinking flexibly, and inhibiting impulses (Diamond, 2013; Ferguson et al., 2021). Multiple studies have found that, to different extents, impairments in executive function are present in various neuropsychiatric conditions such as schizophrenia (Haugen et al., 2021; Raffard et al., 2009; Wobrock et al., 2009), bipolar disorder (Cotrena et al., 2020; Koene et al., 2022), substance abuse (Hester & Garavan, 2004; Kim-Spoon et al., 2017), traumatic brain injury (McDonald et al., 2002), and personality disorders (Gvirts et al., 2012; Koudys & Ruocco, 2022). Importantly, poor executive function performance typically correlates with symptom severity and social maladaptation (Drakopoulos et al., 2020; Simon et al., 2003). However, does executive function perform worse in healthy but socially vulnerable individuals? In other words, does socioeconomically driven chronic stress hinder executive functioning in the absence of a neuropsychiatric condition?

EEG studies have found an association between executive function impairment and alterations in alpha, theta, delta, and beta frequency bands (Bong et al., 2020; Chen et al., 2016). More specifically, researchers have reported an increase in power in slow-wave oscillations such as delta and theta bands during resting state in individuals who have been diagnosed with traumatic brain injury (Dunkley et al., 2015), mild cognitive impairment (Musaeus et al., 2019), Alzheimer’s disease (Babiloni et al., 2020), autism spectrum disorder (J. Wang et al., 2020), and attention deficit hyperactivity disorder (Morillas-Romero et al., 2015), and a decrease in power in such frequency bands in healthy older individuals (Vlahou et al., 2014). Furthermore, cognitive decline in dementias such as Alzheimer’s disease has been associated with an increase in delta-frequency functional connectivity (Babiloni et al., 2004, 2006, 2009; Brunovsky et al., 2003; Cheng et al., 2020; Comi et al., 1998; Locatelli et al., 1998; Meghdadi et al., 2021; Schreiter-Gasser et al., 1994), which has been attributed to an impairment in the neural mechanisms that regulate delta-band coupling. Thus, both slow-wave power and functional connectivity exhibit dynamic changes in the presence of neurological or psychiatric ailments. Are such dynamic changes also present in healthy, but socially vulnerable individuals? If so, are such changes associated with being socially vulnerable, with poorer executive functioning, or with both?

Socially vulnerable populations encompass individuals residing in contexts with limited access to economic resources due to their low-income range (Evans et al., 2005; Evans & English, 2002; Henoch, 2010). These populations are predominantly concentrated in social risk neighbourhoods and lack the necessary social resources to effectively cope with the impacts of external stressors. As a result, their social adaptation (SA) is hindered (Schulte et al., 2022). The cumulative number and intensity of stressors experienced by these individuals, coupled with their inadequate access to stress management resources, contribute to conditions that are conducive to poor mental health and chronic stress (Evans & France, 2022; Evans & Kim, 2010).

In this study, we investigate the EEG spectral and connectivity signatures within a socially vulnerable group, taking into account the unique stressors they face. Specifically, we examine the differences in executive function performance and its association with neural measures between two distinct groups: healthy individuals who are socially vulnerable and those who are socioeconomically stable.

## 2. MATERIALS AND METHODS

### 2.1. Participants

The sample consisted of seventy-six healthy participants, with 38 individuals in the socially vulnerable group and 38 in the control group. Participants’ ages ranged from 34 to 47 years (M = 39.6; [SD = 3.58]), with 43 of them being female. On average, they had 15 years of education (M = 16.0 [3.65]), equivalent to completing primary and secondary education, and no history of psychiatric or neurological conditions. Eligibility criteria for the socially vulnerable group required being a member of a household meeting the 40^th^ percentile qualification for the lowest income range (stretch 1 of 7) of the Chilean Welfare Programme (Ministerio de Desarollo Social y Familia -Gobierno de Chile, 2020). Control participants were recruited from the general population based on accessibility and did not belong to the 40^th^ percentile of the lowest income range.

### 2.2. Data collection

The Social Protection Sheet, issued by the Ministry of Social Development and Family of the Chilean government (Ministerio de Desarollo Social y Familia - Gobierno de Chile, 2020), was used to determine the socioeconomic status of participants in the vulnerable group. This sheet includes comprehensive information on the combined income from labour, pensions, and capital for all household members, as well as the number of individuals within the household and their respective characteristics such as age, disability, or dependency. Additionally, the sheet assesses the access to and ownership of goods and services by the household, enabling an inference of its socioeconomic status based on a comparison with the actual household income received. Only individuals registered within the Social Registry of households were recruited from the vulnerable group.

In accordance with the aforementioned criteria, a brief semi-structured interview was conducted prior to the commencement of the study to assess the level of exposure to long-term social vulnerability for each individual. Only participants who met this criterion were included in the study. Participation was voluntary, and all data were anonymised to ensure confidentiality.

Individuals with visual or hearing impairments, who indicated an inability to complete the assessment battery (e.g., difficulty reading or responding to verbal information or following oral instructions provided by the evaluator), were excluded from participation. Furthermore, individuals with psychiatric or neurological conditions were not included in the study. Prior to their participation, all individuals provided informed consent and signed a consent form. The study received ethical approval from the Adolfo Ibañez University Ethics Committee (Santiago, Chile) and adhered to the protocols outlined in the Declaration of Helsinki.

### 2.3. INECO Frontal Screening (IFS)

All participants completed the IFS, which measures four different executive function components: working memory, motor inhibition, verbal inhibition, and abstraction capacity through eight subtests. The IFS has shown good internal consistency, high reliability, and high concurrent validity (Ihnen et al., 2013; Torralva et al., 2009). IFS performance correlates with other executive function tests too, such as the Frontal Assessment Battery, the Trail Making Test (Part B), the Wisconsin Card Sorting Test, and the verbal phonological fluency test (Baez et al., 2014; Custodio et al., 2016; Gleichgerrcht et al., 2011; Ihnen et al., 2013; Torralva et al., 2009). While the IFS was initially designed for detecting executive dysfunction in dementia, it has proven useful in assessing healthy young and older individuals as well (Fittipaldi et al., 2020; García-Cordero et al., 2017; Sierra Sanjurjo et al., 2019), demonstrating high sensitivity (Moreira et al., 2014; Torralva et al., 2009).

### 2.4. EEG data collection and pre-processing

Participants were sitting during EEG data collection. They were instructed not to think about anything in particular, as often done in resting-state EEG studies (Chennu et al., 2014; Elliott et al., 2005).

We collected a minimum of 10 minutes of resting-state EEG data using a 128-channel high-density EEG system, sampled at 250 Hz and re-referenced to the vertex using a Net Amps 300 amplifier (Electrical Geodesics Inc., USA). EEG data were obtained from both the socially vulnerable group and its matched control group in a state of relaxed wakefulness with eyes open while looking at a fixation cross to minimise eye movement. Eye-blinks were individually assessed to ensure that both groups had their eyes open throughout the recording session.

We measured both eye-blink and eye-movement-related EEG activity. We We derived left and right vertical bipolar electrooculographic (EOG) channels from the raw EEG data by subtracting channels 25 vs 127, and 8 vs 126, respectively (Cologan et al., 2013). Next, we filtered the resulting derived channels with 1-3 Hz to focus on eye-movement-related activity by calculating their standard deviations (SD) using a 1-s non-overlapping sliding window, which was normalised by the mean SD over all windows.

EEG data from 91 scalp channels were selected for further analysis. Channels from non-scalp surfaces such as neck and cheeks were excluded. Continuous EEG data were filtered between 0.5 and 45 Hz and segmented into sixty 10-s epochs. Thus, each epoch was baseline-corrected relative to the mean voltage of the entire epoch. Epochs that contained excessive eye-movement or muscular artefacts were rejected using a quasi-automated procedure whereby abnormally noisy channels and epochs were identified by quantifying their normalised variance – by visual inspection, they were next rejected or kept.

We used independent-component analysis (ICA) using the Infomax ICA algorithm (Bell & Sejnowski, 1995) to identify components and select artefact-related ones. Below 6% of the epochs were rejected. An analysis of variance (ANOVA) revealed no significant difference between the number of epochs selected for each groups. Finally, we interpolated the channels that were rejected by using spherical spline interpolation, and the data were re-referenced to the mean across all channels.

All pre-processing and analysis steps were implemented using MATLAB and the EEGLAB toolbox (Delorme & Makeig, 2004).

### 2.5. Spectral power and phase synchrony analysis

We calculated spectral power within bins of 0.25 Hz by using Fourier decomposition of data epochs and the pwelch method. We converted the power values within the frequency bands *delta* (0–4 Hz), *theta* (4–8 Hz), *alpha* (8–13 Hz), *beta* (13–30 Hz), and *gamma* (30–40 Hz), to relative percentage contributions to the total power. Next, we used cross spectrum between the time-frequency decompositions of each pair of channels to estimate a debiased weighted Phase Lag Index (dwPLI, Vinck et al., 2011).

Phase synchrony is considered a measure of information exchange between neuronal populations and is often calculated from the phase or the imaginary component of the complex cross-spectrum between the signals measured at each pair of channels. For instance, its predecessor, the Phase Locking Value (PLV; Lachaux et al., 1999) is obtained by averaging the exponential magnitude of the imaginary component of the cross-spectrum. However, many of the phase coherence indices derived from EEG data can be affected by differences in volume conduction (Nunez et al., 1997, 1999). As a consequence, a single dipolar source rather than a pair of interacting sources may lead to spurious coherence between spatially disparate EEG channels.

The Phase Lag Index (PLI; Stam et al., 2007) aims at minimising the impact of volume condition and common sources found in the EEG data by averaging the signs of phase differences, thus ignoring average phase differences of 0 or 180 degrees. The rationale behind it is that such phase differences may be likely to be due to volume conduction of single dipolar sources.

Formally, the PLI is defined as the absolute value of the sum of the signs of the imaginary part of the complex cross-spectral density *S*_*xy*_ of two real-valued signals *x*(*t*) and *y*(*t*) at time point or trial *t*:

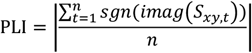

However, PLI has two important limitations: it is very sensitive to noise, and it shows a strong bias towards strong coherences when calculated on small samples. The weighted PLI index (wPLI; Vinck et al., 2011) addresses the former problem by weighting the signs of the imaginary components based on their normalised absolute magnitudes, which yields a dimensionless measure of connectivity unaffected by differences in spectral or cross-spectral power:

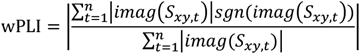

In addition, the dwPLI addresses the latter problem by decreasing its bias when the number of epochs is small. Therefore, we employed the dwPLI measure to estimate functional connectivity.

We quantified the dwPLI peak across all time and frequency bins within each frequency band, for each pair of channels. Thus, we obtained subject-wise and band-wise dwPLI connectivity matrices. Statistical tests were run using Jamovi (The jamovi project, 2020) and JASP (JASP Team, 2023).

## 3. RESULTS

### 3.1. Socially vulnerable and non-vulnerable individuals differ in phase-synchrony connectivity but not in spectral power

To test whether global dwPLI connectivity (i.e., connectivity across all electrodes) differed between groups at any specific frequency band, we entered the dwPLI means into a 4 (frequency band: alpha, beta, theta, delta) × 2 (group: socially vulnerable, control) mixed ANOVA (**Figure 1A**). We found a main effect of frequency band (*F*_(3, 108)_ = 68.87, *p* < .001) and a main effect of group (*F*_(1, 36)_ = 5.64, *p* = .023). Crucially, we found an interaction between these factors (*F*_(3, 108)_ = 3.46, *p* = .019). Holm-Bonferroni-corrected pairwise comparisons revealed a significantly higher dwPLI connectivity in delta (*t*(74) = 4.008, *p* < .001, *d* = 0.92), theta (*t*(74) = 2.54, *p* = .018, *d* = 0.583), and beta (*t*(74) = 2.557, *p* = .018, *d* = 0.587) bands in favour of the socially vulnerable group. We found no significant difference in alpha band (*t*(74) = −0.47, *p* = .639).

**Figure 1.**
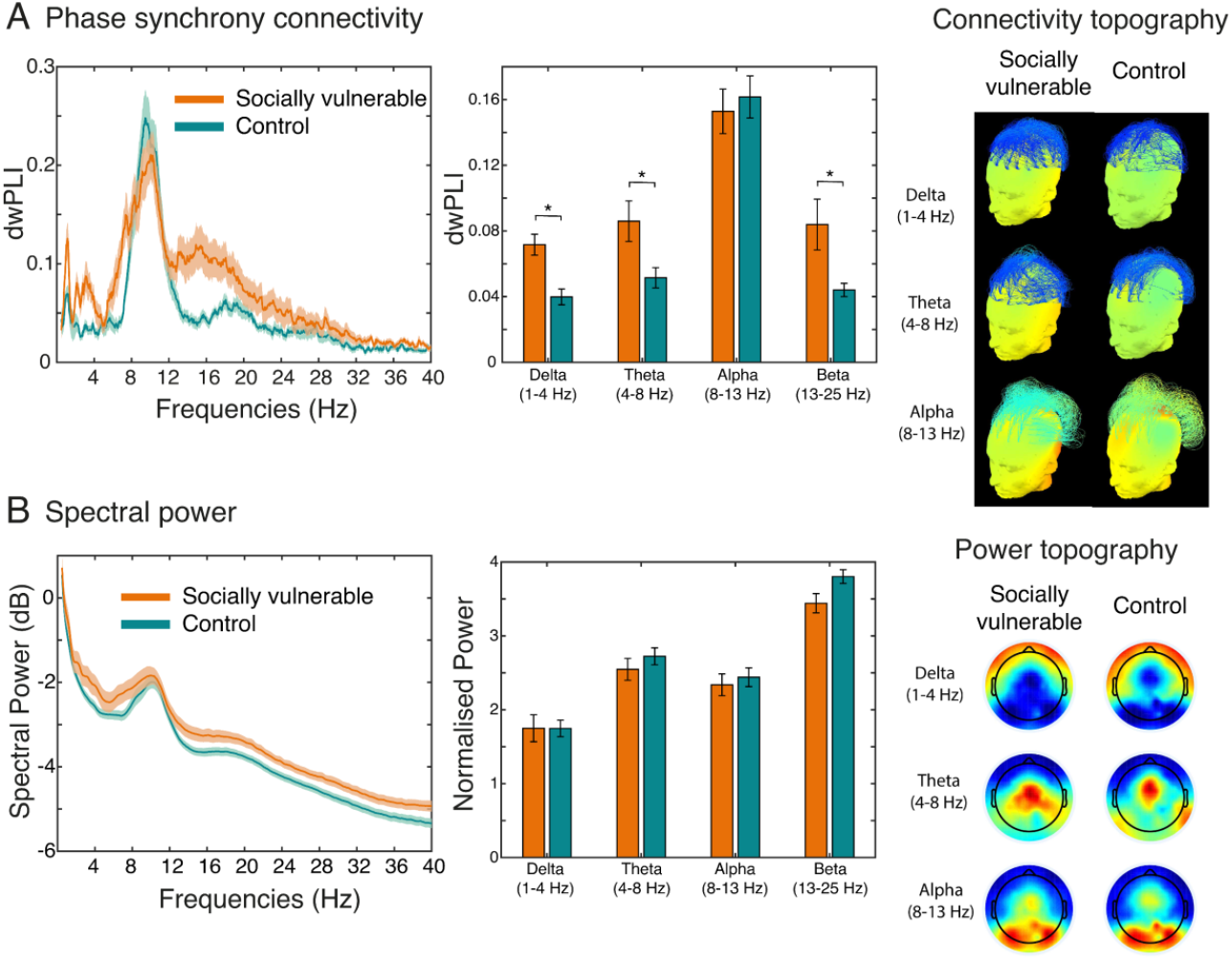
Band-wise phase synchrony connectivity and spectral power in socially vulnerable and control groups. (A) Phase synchrony connectivity. (Left) dwPLI connectivity comparison between groups across frequency bands. (Middle) Differences in dwPLI connectivity between groups: the socially vulnerable group exhibits higher dwPLI connectivity in delta, theta, and beta frequency bands. (Right) Connectivity topographies. Topographic colour maps depicting dwPLI connectivity between pairs of electrodes. (B) Spectral power. (Left) Spectral power comparison between groups across frequency bands. (Middle) We found no spectral power differences between groups. (Right) Power topographies. Topographic colour maps depicting spectral power between pairs of electrodes. Asterisks denote significant differences between conditions. Shaded areas and error bars denote 95% confidence intervals (CI).

Is this increase in phase synchrony connectivity accompanied by an increase in spectral power? To test this, we entered the power values into a 4 (frequency band: alpha, beta, theta, delta) × 2 (group: socially vulnerable, control) mixed ANOVA (**Figure 1B**). We found no main effects or interactions (all p-values > .101).

Together, these results indicate that socially vulnerable individuals exhibit higher phase synchrony connectivity in delta, theta, and beta frequency bands than controls, and that these two groups do not differ in by-frequency-band spectral power.

### 3.2. Executive function and phase synchrony connectivity: bivariate correlations

To test whether executive function correlates with spectral connectivity, we ran a series of exploratory Pearson correlations between IFS scores, dwPLI scores, and demographic data (**Table 1**). We found that delta-(*r* = −0.321, *p* = .006) and theta-band dwPLI connectivity (*r* = −0.241, *p* = .043) negatively correlate with IFS score, irrespective of group, meaning that the lower the IFS score, the higher the dwPLI connectivity score. Next, we found that years of schooling positively correlate with IFS score (*r* = 0.284, *p* = .017) and negatively correlate with dwPLI connectivity in delta (*r* = −0.328, *p* = .004) and theta (*r* = −0.245, *p* = .034) bands, indicating that participants with more years of schooling have lower phase synchrony connectivity in these bands (**Figure 2**).

**Table 1.** Bivariate correlation analysis. Bold and asterisks denote significance: \**p*<.05, \*\**p*<.01, \*\*\**p*<.001.

|  |  | IFS | Delta dwPLI | Theta dwPLI | Alpha dwPLI | Beta dwPLI | Age | Schooling | Gender |
| --- | --- | --- | --- | --- | --- | --- | --- | --- | --- |
| IFS | Pearson's r | — |  |  |  |  |  |  |  |
|  | p-value | — |  |  |  |  |  |  |  |
| Delta dwPLI | Pearson's r | **-0.321\*\*** | — |  |  |  |  |  |  |
|  | p-value | 0.006 | — |  |  |  |  |  |  |
| Theta dwPLI | Pearson's r | **-0.241\*** | **0.483\*\*\*** | — |  |  |  |  |  |
|  | p-value | 0.043 | < .001 | — |  |  |  |  |  |
| Alpha dwPLI | Pearson's r | 0.179 | 0.097 | **0.420\*\*\*** | — |  |  |  |  |
|  | p-value | 0.136 | 0.402 | < .001 | — |  |  |  |  |
| Beta dwPLI | Pearson's r | -0.031 | **0.442\*\*\*** | **0.575\*\*\*** | **0.375\*\*\*** | — |  |  |  |
|  | p-value | 0.800 | < .001 | < .001 | < .001 | — |  |  |  |
| Age | Pearson's r | -0.229 | 0.106 | 0.206 | 0.144 | 0.163 | — |  |  |
|  | p-value | 0.053 | 0.364 | 0.074 | 0.216 | 0.160 | — |  |  |
| Schooling | Pearson's r | **0.284\*** | **-0.328\*\*** | **-0.245\*** | -0.005 | -0.163 | -0.081 | — |  |
|  | p-value | 0.017 | 0.004 | 0.034 | 0.963 | 0.162 | 0.486 | — |  |
| Gender | Pearson's r | -0.212 | 0.085 | -0.067 | -0.047 | -0.118 | 0.181 | -0.067 | — |
|  | p-value | 0.073 | 0.467 | 0.566 | 0.689 | 0.310 | 0.115 | 0.565 | — |

**Figure 2.**
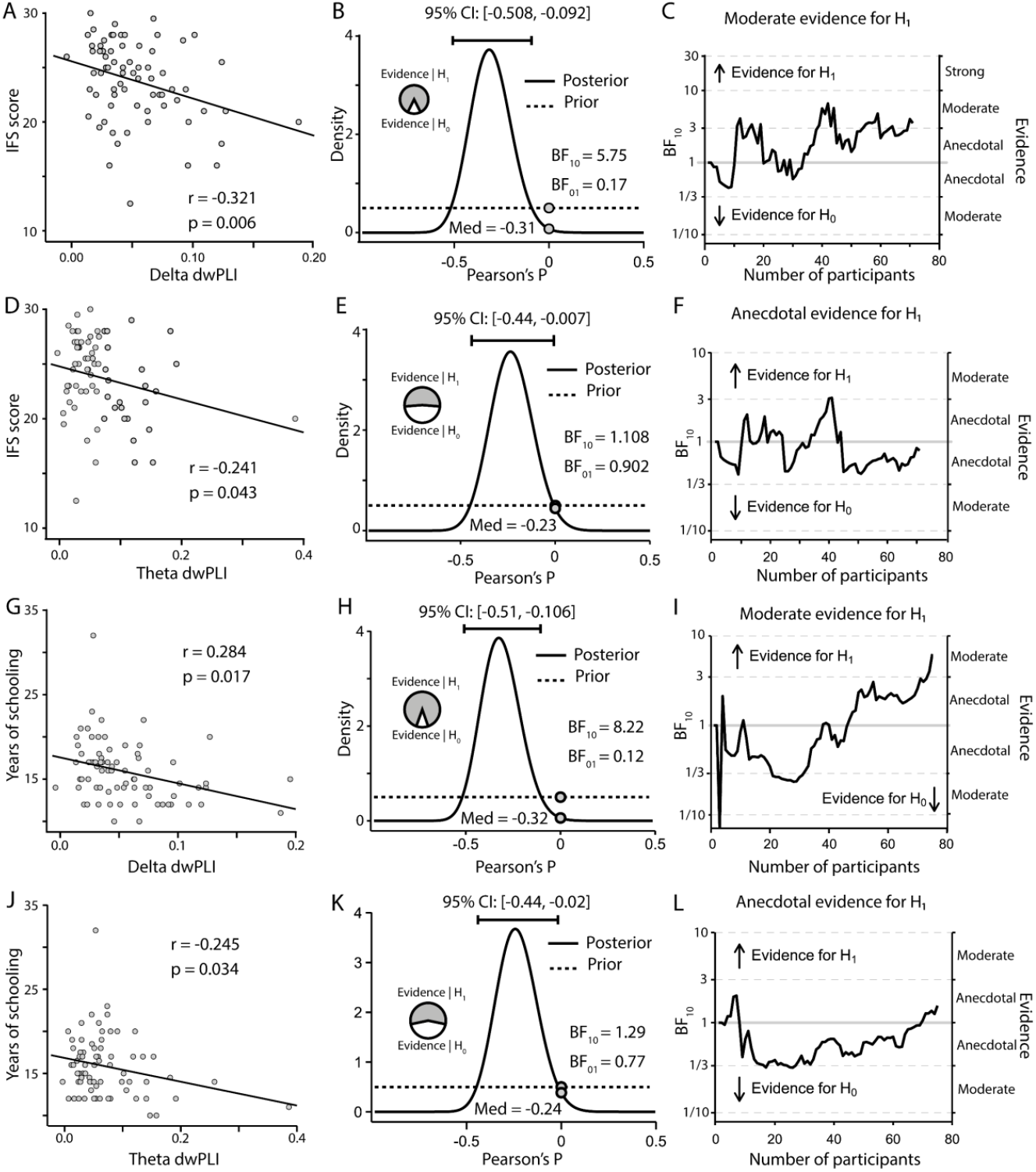
Scatterplots and Bayesian assessment of the evidence. (A-C) IFS score and delta-band dwPLI. (A) IFS score and delta dwPLI showed a significant negative correlation. (B) Bayes factors provided moderate evidence in favour of the alternative hypothesis model (i.e., IFS score and delta dwPLI correlate), which is depicted by the estimated population effect size, with a median of -0.31 and a 95% central credible interval of -0.508 and -0.092. (C) Sequential analysis shows that most participants give moderate or anecdotal support to the alternative hypothesis model. (D-F) IFS score and theta-band dwPLI. (D) IFS score and theta dwPLI showed a significant negative correlation. (E) Bayes factors provided anecdotal evidence in favour of the alternative hypothesis model (i.e., IFS score and theta dwPLI correlate), which is depicted by the estimated population effect size, with a median of -0.23 and a 95% central credible interval of -0.44 and -0.007. (F) Sequential analysis shows that all participants give anecdotal support either in favour or against the alternative hypothesis model. (G-I) Years of schooling and delta-band dwPLI. (G) Years of schooling and delta dwPLI showed a significant negative correlation. (H) Bayes factors provided moderate evidence in favour of the alternative hypothesis model (i.e., years of schooling and delta dwPLI correlate), which is depicted by the estimated population effect size, with a median of -0.32 and a 95% central credible interval of -0.51 and -0.106. (I) Sequential analysis shows that most participants give moderate or anecdotal support to the alternative hypothesis model. (J-L) Years of schooling and theta-band dwPLI. (J) Years of schooling and theta dwPLI showed a significant negative correlation. (K) Bayes factors provided anecdotal evidence in favour of the alternative hypothesis model (i.e., years of schooling and theta dwPLI correlate), which is depicted by the estimated population effect size, with a median of -0.24 and a 95% central credible interval of -0.44 and -0.02. (L) Sequential analysis shows that most participants give anecdotal support to the null hypothesis model.

### 3.3. Controlling for age, schooling, and gender further supports differences between groups in executive function and delta phase-synchrony connectivity

#### 3.3.1. IFS performance

To further explore the effects of group and phase synchrony connectivity, we ran an ANCOVA whereby age, years of schooling, and gender were included as covariates. First, we further explored the difference in IFS score between groups (**Figure 3A**). We found again that the socially vulnerable group (M = 22.5 [CI = 21.2, 23.8]) performed worse at the IFS than the control group (25[23.7, 26.2]), (*F*_(1, 65)_ = 5.613, *p* = .021). We did not find any other significant effects (all p-values > .144).

**Figure 3.**
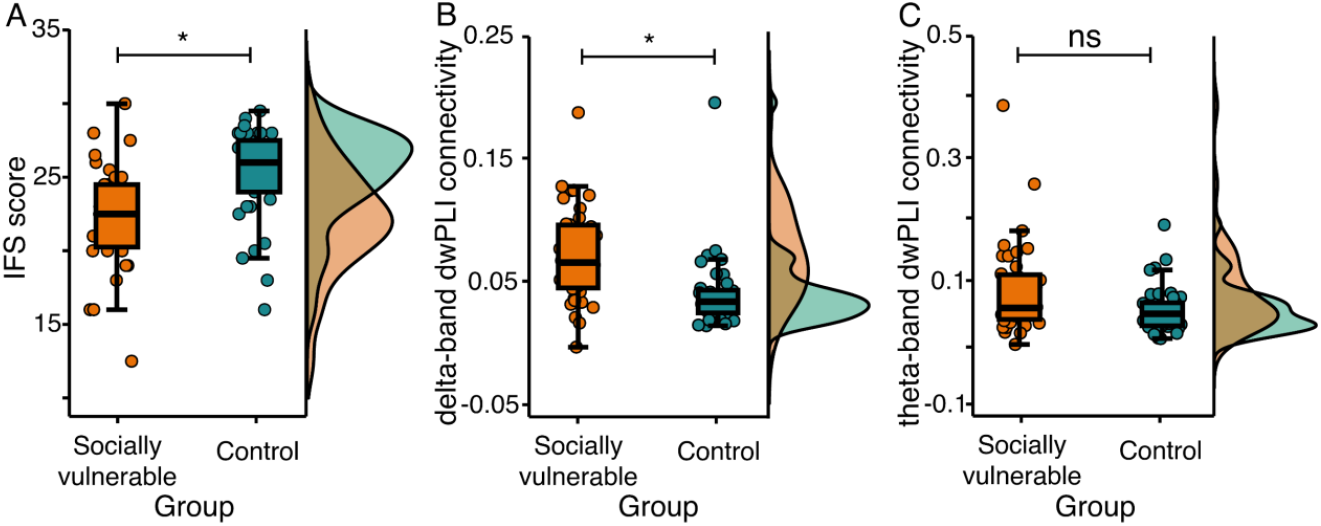
Group differences based on the ANCOVA model. (A) IFS scores were significantly lower in the socially vulnerable group compared to the control group. (B) delta-band phase synchrony connectivity was significantly higher in the socially vulnerable group compared to the control group. (C) theta-band phase synchrony connectivity did not significantly differ between groups. Asterisks denote significant differences between groups. Error bars denote 95% CI.

#### 3.3.2. Phase synchrony connectivity

Next, we ran a series of ANCOVA models to test the effects on dwPLI connectivity for each frequency band, separately. The ANCOVA model for delta-band connectivity (**Figure 3B**) showed higher dwPLI for the socially vulnerable group (0.067[0.054, 0.08]) compared to the control group (0.042[0.03, 0.06]), (*F*_(1, 69)_ = 5.77, *p* = .019). We did not find any other significant effects (all p-values > .263).

The ANCOVA model for theta-band dwPLI connectivity (**Figure 3C**) did not find an effect of group (*F*_(1, 69)_ = 1.28, *p* = .261). We did not find other significant effects (all p-values > .108). Exploratorily, we took out years of schooling from the ANCOVA model and found differences between groups (*F*_(1, 71)_ = 4.96, *p* = .029), which may suggest that theta-band connectivity is moderated by schooling (see further below).

### 3.4. Delta-band synchrony and years of schooling predict membership in either the vulnerable or control groups

To determine the likelihood of belonging to either the vulnerable or control group, a binomial logistic regression was conducted, with several variables of interest serving as predictors (*F*_(1, 71)_ = 4.96, *p* = .029), including IFS score, delta-band dwPLI, theta-band dwPLI, years of schooling, and age. This regression model enabled us to statistically predict which cognitive, demographic, and neural parameters could effectively classify participants into their respective groups and with what degree of accuracy. The model provided an explanation for 54% (R^2^_CS_) to 72% (R^2^_N_) of the variance in the dependent variable, resulting in a classification precision of 86%, exhibiting a sensitivity of 82% and a specificity of 89% (**Table 2**). Notably, two predictors significantly contributed to this model: delta-band dwPLI (*X*^2^_(1)_ = 12.39, *p* < .001) and years of schooling (*X*^2^_(1)_ = 22.21, *p* < .001), (**Figure 4**).

**Table 2.** Binomial logistic regression model coefficients. Estimates represent the log odds of Socially vulnerable group vs. Control group. Estimates represent log odds of “Group = Socially vulnerable” vs. “Group = Control”. Bold and asterisks denote significance: *p<.05, **p<.01, ***p<.001

| Predictor | Estimate | 95% Confidence Interval |  | SE | Z | p | Odds ratio | 95% Confidence Interval |  |
| --- | --- | --- | --- | --- | --- | --- | --- | --- | --- |
|  |  | Lower | Upper |  |  |  |  | Lower | Upper |
| Constant | 13.55511 | 0.0634 | 27.0468 | 6.884 | 1.9692 | 0.049 | 770743.573 | 1.0655 | 5.58e+11 |
| IFS Score | -0.15251 | -0.3971 | 0.0920 | 0.125 | -1.2224 | 0.222 | 0.859 | 0.6723 | 1.096 |
| delta dwPLI | 55.71430 | 17.1421 | 94.2865 | 19.680 | 2.8310 | **0.005\*** | 1.57e+24 | 2.78e +7 | 8.87e+40 |
| Gender: |  |  |  |  |  |  |  |  |  |
| Female – Male | -1.06581 | -2.8191 | 0.6875 | 0.895 | -1.1914 | 0.233 | 0.344 | 0.0597 | 1.989 |
| Schooling | -0.74251 | -1.1773 | -0.3078 | 0.222 | -3.3474 | **<.001\*** | 0.476 | 0.3081 | 0.735 |
| Age | 0.00379 | -0.2224 | 0.2300 | 0.115 | 0.0328 | 0.974 | 1.004 | 0.8006 | 1.259 |
| theta dwPLI | -8.29594 | -28.4988 | 11.9070 | 10.308 | -0.8048 | 0.421 | 2.50e -4 | 4.20e-13 | 148294.659 |

**Figure 4.**
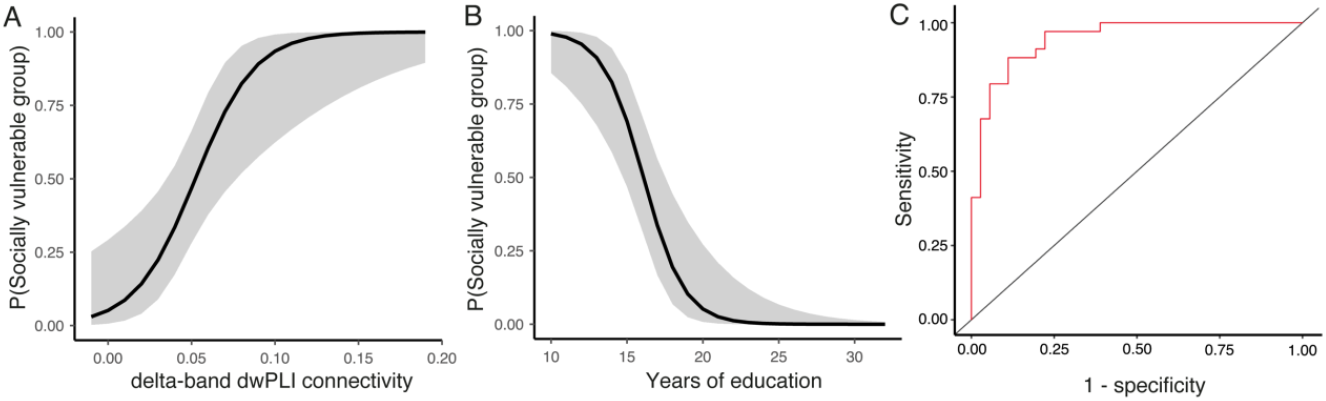
Binomial logistic regression model. (A) Performance of delta-band dwPLI connectivity as predictor of group classification. (B) Performance of years of schooling as predictor of group classification. (C) Receiver Operating Characteristic (ROC) curve represents the classification performance of the model (AUC = 0.94).

Together, these results indicate that years of schooling and delta-band dwPLI connectivity contribute the most to predicting an individual’s group (i.e., socially vulnerable or control).

## 4. DISCUSSION

Socially vulnerable individuals often experience chronic stress due to limited access to education, healthcare, safe environments, and work opportunities, which poses a threat to their cognitive development and mental health (Cermakova et al., 2018; De Nadai et al., 2020; Engelberg et al., 2016; Giles-Corti & Donovan, 2002; Migeot et al., 2022). In this study, we investigated how social vulnerability affects executive functioning and its relationship with resting-state neural activity in neural power frequency and connectivity spaces in healthy individuals. Our main findings revealed that socially vulnerable individuals display higher delta, theta, and beta functional connectivity compared to controls. Additionally, both delta- and theta-band functional connectivity exhibited a negative association with executive functioning, indicating that connectivity increases as executive functioning scores decrease. Interestingly, there was no evidence of neural differences in power bands between the two groups. Finally, after accounting for covariates such as age, years of schooling, and gender, we found that only the increase in delta-band connectivity predicted social vulnerability, while theta-band connectivity was associated with years of schooling.

Prior studies have reported an association between both higher resting-state delta-band power and connectivity, and cognitive decline in conditions such as mild cognitive impairment and dementia (Adler et al., 2003; Babiloni et al., 2009; Brunovsky et al., 2003; Kwak, 2006; Locatelli et al., 1998). Although findings in this area are still inconclusive (C. Wang et al., 2022), an association between slow-wave connectivity and cognitive performance has been established (Laptinskaya et al., 2020). Interestingly, while it is common to come across studies that compare cognitively impaired patients with healthy controls, demonstrating differences in delta-band power and other bands (Torres-Simón et al., 2022), the focus has understandably shifted towards examining neural networks and connectivity measures (Liu et al., 2023) as brain connectivity is perceived as more interpretable in a context of the brain as a complex neural network. In our study, we did not find evidence for power differences in any frequency band between socially vulnerable individuals and controls. However, we found connectivity differences between groups and in association with executive function performance, particularly for slow frequencies. It is important to note that suspected cognitive differences observed in pre-dementia studies and socioeconomic status studies are unlikely to be similar, and the same expectation applies to neural markers. Our findings suggests that the mechanisms by which neurological disorders and socioeconomic factors impact executive functioning are different. On the other hand, based on behavioural evidence, Mani et al. (2013) found that individuals facing poverty or resource scarcity exhibit a diminished capacity to make rational decisions and solve complex problems. Persistent concerns about economic matters were observed to adversely affect cognitive abilities, resulting in lower performance on cognitive tasks and a decline in executive function. In other words, the constant preoccupation with resource scarcity may negatively impact brain function, similar to other factors that deplete cognitive capacity, such as sleep deprivation or alcohol consumption. The aforementioned findings support the notion that socioeconomic vulnerability, such as poverty, is associated with lower cognitive performance, given that economic concerns consume cognitive resources. This hypothetical effect could be mirrored at the neural level by a higher predominance of connectivity in low-frequency bands.

Our findings indicate that the increased delta-band connectivity observed in the socially vulnerable group cannot be accounted for by differences in age, years of schooling, gender, or IFS score – only group membership significantly predicted changes in delta-band connectivity. This raises the question of which specific underlying factor, present among socially vulnerable individuals, drives this effect. While years of schooling may seem like a straightforward factor to predict group membership, as economic and social hardships often lead to school dropout (Adelman & Székely, 2017; Zaff et al., 2017), it is noteworthy that delta-band dwPLI connectivity contributed the most to predicting an individual’s group (socially vulnerable or control). This leaves the door open for further research to explore its functional role and potential involvement in the underlying mechanisms of neural reconfiguration in response to hardship.

Recent studies have brought attention to the early effects of socioeconomic hardship on neural development, specifically in relation to neural markers of power in EEG. For instance, Wilkinson et al. (2023) found associations between family income and neural power in a USA sample, whereas Otero (1997) obtained similar findings in a Mexican sample. Interestingly, while these findings support the early effects of hardship on neural development, the evidence for adults is not as conclusive. This suggests that while neural power may capture the initial effects of hardship, other neural signatures likely come into play later in life.

In a recent study, we found that cognitive variables (working memory and fluid intelligence) and socio-affective variables (self-esteem, stress, and locus of control) can predict social adaptation among adults living in vulnerable contexts (Neely-Prado et al., 2019). Specifically, we found that 31.8% of the differences in social adaptation were accounted for by stress, internal locus of control, and self-esteem, while 7% depended on working memory and fluid intelligence. The current study aimed to explore the relationship between neural markers and executive functioning in both socially vulnerable and non-vulnerable groups. Interestingly, some of the results indicate changes in slow-wave connectivity, which may also be related to or underlie socio-affective aspects of cognition. The combined findings of these two studies suggest that public policies could target self-esteem, locus of control, and perceived stress as relevant areas for intervention. Additionally, neural markers may play a role in defining the degree of belonging and could potentially be used to track progress and understand the underlying neural mechanisms of vulnerability in socio-economic hardship.

Our study has one limitation: we did not measure any physiological marker of stress. Although there is well-established evidence that socially vulnerable individuals experience higher stress levels due to socioeconomic factors (Cermakova et al., 2018; Migeot et al., 2022), this omission hinders our ability to delve deeper into the relationship between IFS, neural connectivity, and chronic stress. However, it is worth noting that our team collaborated with the same population as the participants recruited in this laboratory study during the fieldwork conducted by Neely-Prado et al. (2019). During that collaboration, it was observed that the perception of stress in individuals from socially vulnerable backgrounds emerged as a significant characteristic. Future studies should collect data on physiological markers of stress, allowing for a more comprehensive exploration of statistical nuances that our current analyses may have missed. Moreover, future studies should also investigate the specific executive functions most impacted by social vulnerability and how they evolve throughout childhood and adolescence. Understanding how slow-wave connectivity develops during the cognitive development of socially vulnerable individuals may shed light on how social factors affect cognitive development and the emergence of susceptibility to neuropsychiatric disorders among this group.

In summary, our study explored the impact of social vulnerability on executive functioning and its association with neural connectivity in healthy individuals, and found that socially vulnerable individuals exhibit higher slow-wave neural connectivity than non-vulnerable individuals.

## 6. ACKNOWLEDGEMENTS

David Huepe is supported by an ANID/FONDECYT Regular (1231117) research grant. The content of this article is solely the responsibility of the authors and does not represent the official views of this institution.

